# High speed, three-dimensional imaging reveals chemotactic behavior specific to human-infective *Leishmania* parasites

**DOI:** 10.1101/2020.07.30.220541

**Authors:** Rachel C. Findlay, Mohamed Osman, Kirsten A. Spence, Paul M. Kaye, Pegine B. Walrad, Laurence G. Wilson

## Abstract

Cellular motility is an ancient eukaryotic trait, ubiquitous across phyla with roles in predator avoidance, resource access and competition. Flagellar-dependent motility is seen in a variety of parasitic protozoans and morphological changes in flagellar structure and function have been qualitatively described during differentiation. However, whether the dynamics of flagellar motion vary across lifecycle stages and whether such changes serve to facilitate human infection is not known. Here we used holographic video microscopy to study the pattern of motility in insect midgut forms of *Leishmania* (procyclic promastigotes; PCF) and differentiated human infective metacyclic promastigotes (META). We discovered that PCF swim in a slow, corkscrew motion around a gently curving axis while META display ‘run and tumble’ behaviour in the absence of stimulus, reminiscent of bacterial behaviour. In addition, we demonstrate that META specifically respond to a macrophage-derived stimulus, modifying swimming direction and speed to target host immune cells. Thus, the motility strategy employed by *Leishmania* appears as a random search that is replaced with a ballistic swimming motion in the presence of an immunological stimulus. These findings shed unique insights into how flagellar motion adapts to the particular needs of the parasite at different times in its lifecycle and define a new pre-adaptation for infection of the human host.

---

Flagellar-dependent motility [1, 2] is key for transmission of unicellular *Leishmania* parasites, causative agent of the leishmaniases. These infections represent the world’s ninth largest infectious disease burden and threaten 350 million people globally [3]. Like many protozoan parasites, life-cycle stage-specific differentiation is observed, affecting both *Leishmania* body shape and flagellar morphology [4, 5, 6]. Yet *Leishmania* parasites are unusual among flagellated microorganisms in the drastic extent the flagella length changes between stages. Procyclic promastigotes (PCF) cell bodies are 10-12 *μ*m long, with a flagellum of approximately the same length, while human-infective metacyclic promastigotes (META) cell bodies are 8-10 *μ*m in length, with a flagellum 20 *μ*m long [7].

Studying the three-dimensional swimming patterns of motile flagellates gives insight into regulation of the flagellar beat and details of the cells’ navigation strategy. The physics of microorganism swimming and navigation has been considered extensively in the context of bacterial motility, where biased random walks are used to counter the randomisation of Brownian motion [8, 9, 10]. *Leishmania* cells are an order of magnitude larger in each dimension than typical model bacteria, and rotational diffusivity scales with the cell’s volume. In a medium with viscosity close to that of water, it takes a few seconds to randomise the orientation of an *E. coli* cell, but over 200 seconds to randomise the orientation of *Leishmania*. The physical constraints that shape the response of these two micoorganisms are therefore fundamentally different. Both live in a low-Reynolds number environment from a fluid dynamics perspective, but Brownian motion dominates life for the bacterial system, while it is negligible for these eukaryotes. Nevertheless, there are intriguing signs that the run-tumble locomotion characteristic of bacteria likely *E. coli* has analogues in motile single-cell eukaryotes: *Chlamydomonas reinhardtii* algae exhibit run and tumble behaviour[11], and sperm from the *Arbacia punctulata* sea urchin exhibit sharp reorientation events [12].

Holographic microscopy has proven to be a versatile and powerful tool for investigating the dynamics of swimming microorganisms at high speed and diffraction-limited resolution [13, 14, 15, 16, 12]. Among the advantages of three-dimensional tracking is the ability to unambiguously resolve chirality in the shape of objects and in the geometry of swimming paths. A long-standing question in the biological physics of the organelle at the heart of the axoneme focuses on the breaking of symmetry. The axoneme has a chiral structure in which dynein molecules on each doublet can attach to their clockwise neighbour, as viewed from the basal end of the axoneme [17]. This rotational symmetry is broken by other mechanical and beat regulation processes to allow both left- and right-handed beats in *Plasmodium* microgametes [15]. Here we investigate and define the swimming dynamics and presence of a chemotactic strategy in *Leishmania mexicana*, and examine the role of chirality in this species’ swimming behavior.

## Results and Discussion

We generated PCF and META stages of *Leishmania mexicana* using molecularly-verified culture techniques and confirmed the purity of our populations by assessing morphological characteristics and measuring abundance of mRNA for two lifecycle stage-defining markers, *Histone H4* and *SHERP* [18, 19] (Fig. 1a). We then used holographic microscopy to analyse the three-dimensional motility of isolated parasites. Subsets of this data are presented as composite renderings of five hundred cell tracks in a sample volume ∼ 1.5 × 1.5 × 1.2 mm^3^ (Fig. 1b,c). In liquid culture, observed swimming speeds vary both within and between these two groups.

**Figure 1:**
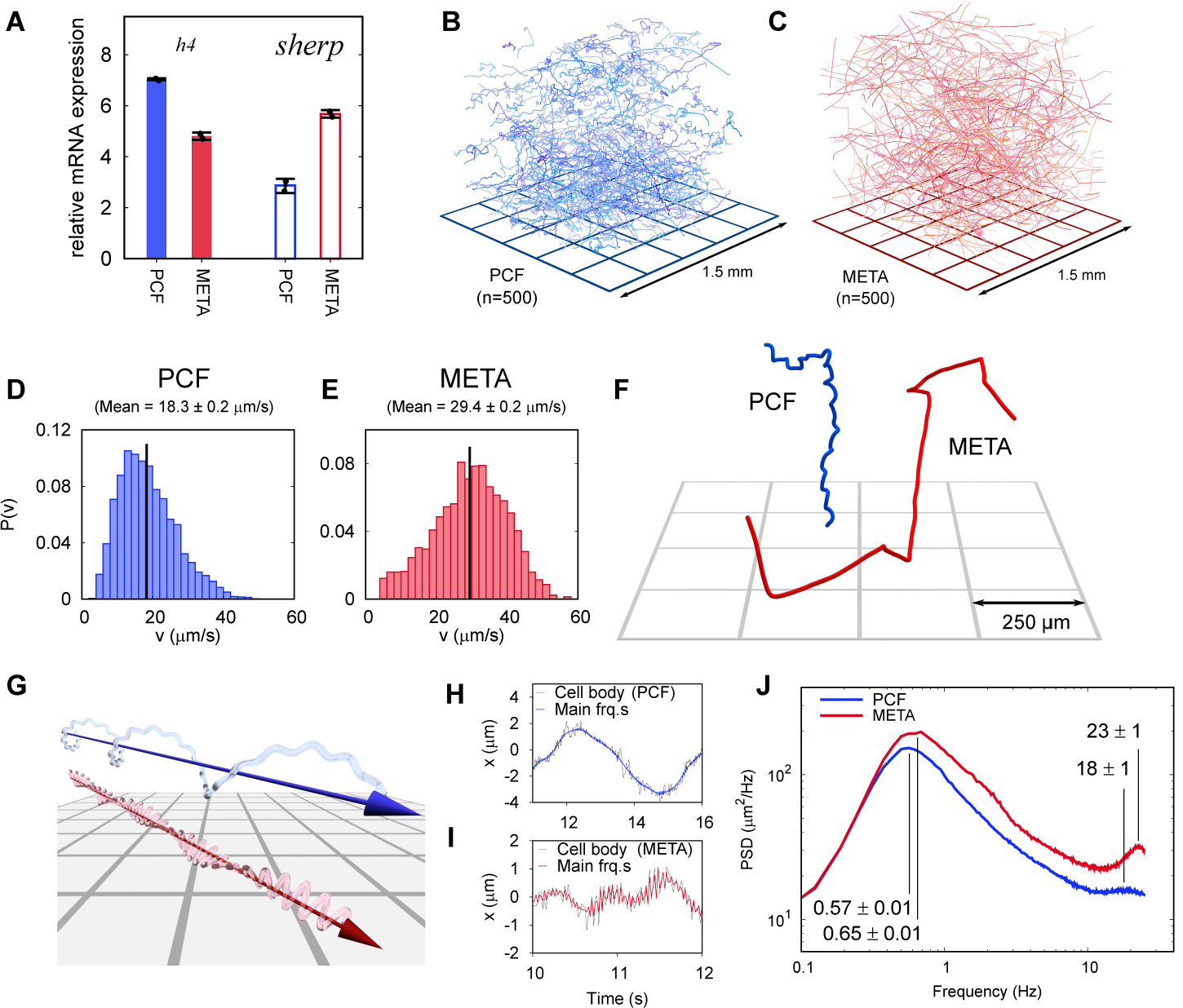
Swimming phenotypes in infective and non-infective *L. mexicana* promastigotes. (A), qPCRs of *H4* (PCF molecular marker) and *SHERP* (META molecular marker) levels relative to *N-myristyltransferase* (*nmt*) (constant transcript control) demonstrate that enriched, but not wholly distinct populations of cells were isolated and characterised. (B), Three-dimensional swimming trajectories from 500 PCF cells of *L. mexicana*, showing stereotypical meandering helical paths. (C), Trajectories of 500 META cells of *L. mexicana*. These show a marked transition to a ‘run and tumble’ phenotype of rapid, straight trajectories interspersed by sharp reorientations. (D,E), Distribution of instantaneous swimming speeds in PCF (n = 3231) and META (n = 2202) cells, respectively. (F), Individual tracks of PCF and META cells at a smaller scale, illustrating the different swimming phenotypes. (G), Cartoon illustrating the dominant swimming phenotype for PCF (top) and META (bottom). (H), The x-component of cell body motion as a function of time for a PCF cell. The black line represents the raw data, and the blue line is the motion reconstituted using the principal frequency components only. (I), x-component of motion for a META cell, both raw data (black) and principal frequencies only (red). The flagellar beat frequency is more pronounced in META cells compared to PCF cells because their cell bodies of the former are significantly smaller, and their flagella are longer. (J), Averaged power spectra of PCF (blue, n = 3231) and META (red, n = 2202) motion, when the lowest frequencies have been removed (see text). The peaks at around 0.6 Hz correspond to the rotation of the flagellar beat plane and the peak at around 20 Hz to the flagellar beat in both cell types. The peak values and uncertainties are indicated.

PCF cells (Fig. 1d) swim significantly slower, with a mean of 18.3 ± 0.2 *μ*m/s (SEM), while META cells (Fig. 1e) have a mean speed of 29.4 ± 0.2 *μ*m/s (SEM). The speed overlap between these populations may be due to natural variability, though we cannot rule out minor levels of contamination during purification (Fig. 1a). Our results indicate PCF cells swim with a characteristic slow, corkscrew-like motion around a gently curving axis, a similar pattern to that of sea urchin sperm cells [12]. In contrast, culture-derived META cells display a distinct swimming phenotype with straight path segments punctuated by sharp turning events, a ‘run and tumble’ motif reminiscent of enteric bacteria such as *Escherichia coli* (Fig. 1b). This analogy should be applied advisedly, however as bacterial runs and tumbles are a strategy to enable navigation in an environment that randomises their swimming direction through rotational Brownian motion. *Leishmania* cells are at least 5-8 times the (linear) size of *E. coli* bacteria so are less impacted by Brownian motion. As rotational diffusivity is proportional to particle volume, *Leishmania* will lose their orientation around 100 times more slowly than *E. coli* [8]. It therefore seems likely that the run and tumble behaviour observed here originates from a different imperative serving a distinct function specific to *Leishmania*. We note that foraging strategies among larger animals follow superficially similar movement patterns [20].

While holography under high magnification can reveal the geometry of eukaryotic flagella [15, 21], the flagellar beat frequency can be inferred even from low-magnification (i.e. large-scale) tracking data. The movement of the *Leishmania* cell body carries a residual signature of the flagellar motion, and this is apparent when the smoothed swimming trajectories (e.g. Fig. 1f) are subtracted from the raw tracking data. Examples of this residual motion are shown for PCF (Fig. 1h) and META (Fig. 1i). The frequency-space representations of these data show intensity peaks at around 0.5 and 20 Hz, which correspond to rotation of the flagellar beat plane (‘body roll’) and the flagellar beat frequency, respectively (see methods for details). The distinct swimming phenotypes of PCF and META cells are illustrated in Fig. 1g. Applying a frequency-domain filter that excludes information outside these principal frequency bands results in the blue (PCF) and red (META) curves in Figs. 1h and 1i, demonstrating that the body roll and flagellar beat are largely responsible for the residual signal. Figure 1j shows the power spectrum of the residual cell motion averaged over populations of PCF and META cells. The beat plane rotation frequency is approximately the same in both cases: 0.57 ± 0.01 Hz and 0.65 ± 0.01 Hz for PCF and META cells, respectively. In sharp contrast, the flagellar beat frequency is significantly higher in the case of the metacyclic cells 2±1 Hz versus 18±1 Hz for procyclic cells. These results show how faster swimming is achieved after differentiation: META cells have a flagellar beat that is around 27% faster than the PCF cells, but with a beat plane that rotates only 15% faster, along with an average swimming speed that is over 60% faster than PCF cells. Current understanding is that the interplay of Dynein activation and microtubule stiffness determine the beat frequency [22], suggesting that the internal structure of the flagellum is remodelled in the transition from PCF to META.

The chirality of swimming motion can be challenging to access experimentally [14, 23, 24], especially in cells with a complicated morphology. We assess the chirality of the tracks by using three-dimensional tracking to measuring the helicity *H* [15] as illustrated in Fig. 2a. Importantly, neither PCF (Fig. 2b) nor META (Fig 2c) cell tracks display a systematic chirality on the population level. Despite this, individual tracks do display a right- or left-handed character, which we speculate is a consequence of small asymmetries in individual cell bodies, or flagellar attachment points.

**Figure 2:**
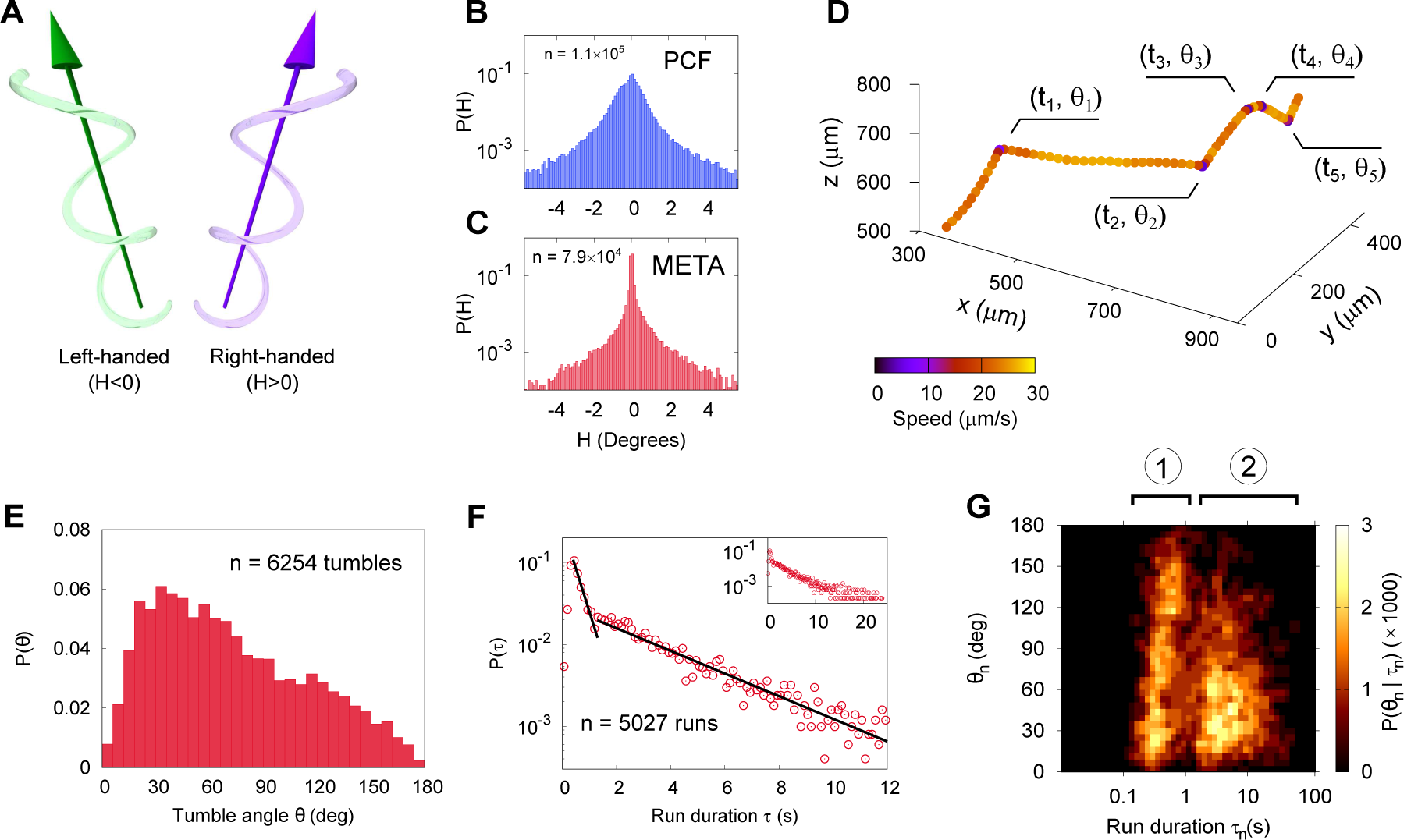
Three-dimensional geometry of META and PRO cell tracks in *L. mexicana*. (A), Cartoon depicting a left-handed track (helicity H¡0) and a right-handed track (H¿0). The arrows indicate the large-scale swimming displacement. Helicity is defined such that H=0 corresponds to a straight line. (B), The normalised probability density of instantaneous helicities in a population of PCF (n = 1.1×10^5^). Individual cells tend to have left- or right-handed characteristic swimming trajectories, but whole populations do not exhibit handedness and are remarkably balanced. These data are drawn from all cells at all times; distributions of helicity from individual cells; helicity per cell is shown in Supplementary Materials. (C), Normalised probability density of helicity among META (n = 7.9×10^4^). These cells display a much narrower range of helicities, commensurate with the straighter trajectories, while also retaining the equal distribution of left- and right-handed instances. (D), An example META trajectory in which the colour code shows instantaneous speed. The cell was imaged at 50Hz, with every 30th point plotted for clarity. Five changes in direction (‘tumbles’) are identified in which the cell’s orientation changes through an angle n, at time *tn*. We define run durations as the delay time between two tumbles *τ*_n_ = *t*_n_ − *t*_n−1_. (E), the normalised probability density of tumble angles. Many tumbles change the swimming direction through an angle of less than 90^°^, with a mean tumble angle of 72^°^± 0.5^°^ (SEM) and a mode of 21^°^. (F), The distribution of run durations. These show two distinct exponential regimes, at short and long times as indicated by the piecewise straight-line fits (black lines). The inset shows the complete range of run durations. Data are noisy at ¿ 10 seconds due to the limited size of the sample volume. (G), Correlation between the duration of a run and the tumble angle following it (n = 5,027 runs). Shorter runs (population 1) are followed by a wider range of angles than longer runs (population 2) indicative of two characteristic run phenotypes.

To analyse the characteristic run and tumble nature of META tracks in more detail, we separate them into ‘runs’ of duration *τ*_n_ = *t*_n_ − *t*_n−1_ punctuated by ‘tumbles’ through an angle *θ* (Fig. 2d). Tumbles appear biased towards small angles (Fig. 2e), so several tumbles would typically be required to completely randomise a cell’s swimming direction. In contrast to commonly studied bacterial species that display a heuristically similar swimming pattern, the distribution of *Leishmania* META run durations is not a simple exponential. We observe at least two distinct run behavioural ‘regimes’, as shown in Fig. 2f. Runs of *τ* ¡ 1.5 seconds are the most common, though the distribution has a long exponential tail. The two distinct run regimes are denoted by exponential lines of best fit (non-linear least squares regressions over the domain as indicated) in the main panel of Fig. 2f, while the inset shows all data down to very low event frequencies at times up to 25 seconds. Figure 2g shows the quantified relationship between a run duration (*τ*) and the subsequent tumble angle (*θ*). Shorter runs are followed by a broad range of tumble angles (population 1), while longer runs are followed by a smaller angle tumble which preserves the cell’s swimming direction (population 2).

In bacterial systems [8, 25], an exponential distribution of runs is the hallmark of an underlying Poisson process with a constant probability of tumbling per unit time. Fine tuning of this tumble probability in response to a biological sensing mechanism enables cells to navigate toward or away from an external stimulus. To determine whether the behaviour of META cells was fixed or could be regulated in conditions that might favour intracellular infection, we repeated our analysis using META cells in the presence of both murine and human macrophages, with results shown in Fig. 3. META parasites were exposed to medium alone versus J774.2 immortalised, mouse stomach-derived macrophages (mM*ϕ*) and human blood-derived macrophages (hM*ϕ*). We compared the swimming behaviour of META cell populations close to the chemotactic bait (BAIT; ‘Tip’) with those in a region 1 cm from the bait (‘Away’), with results shown in Fig. 3a. The cells were significantly drawn toward the BAIT when macrophages are present, with the primary human blood-derived macrophages a stronger draw as expected given their greater biological relevance. The number of tumbles decreases threefold in the presence of hM*ϕ* (Fig. 3b), independent of the number of tracks observed or track length. A more detailed examination of the geometry of the tracks (quantities recapitulated in Fig. 3c) yields unaltered distributions of tumble angles (Fig. 3d) and run durations (Fig. 3e) between control and hM*ϕ* response. The average displacement per run is enhanced in both ‘Tip’ and ‘Away’ cases in the presence of hM*ϕ* (Fig. 3f). This is consistent with a small population of cells retaining the default run-tumble behaviour, while the majority of cells swim faster displaying far fewer, or no tumbles. Importantly, this indicates the presence of a biologically-relevant chemotactic stimulus from hM*ϕ* that causes META cells to swim faster and straighter. Speed distributions for each experimental condition are shown in Fig. 3g-l. There is no distinction between the ‘tip’ and ‘away’ speed in the presence of DMEM alone. In contrast, the presence of mM*ϕ* enriches the fraction of cells swimming at low speeds (¡10 *μ*m/s), while in the presence of hM*ϕ* the average speed is increased by 30%. These distinct phenotypic responses highlight the specificity of the parasite response, and suggest the presence of a soluble, macrophage-derived stimulus to which human-infective META promastigotes are highly sensitive.

**Figure 3:**
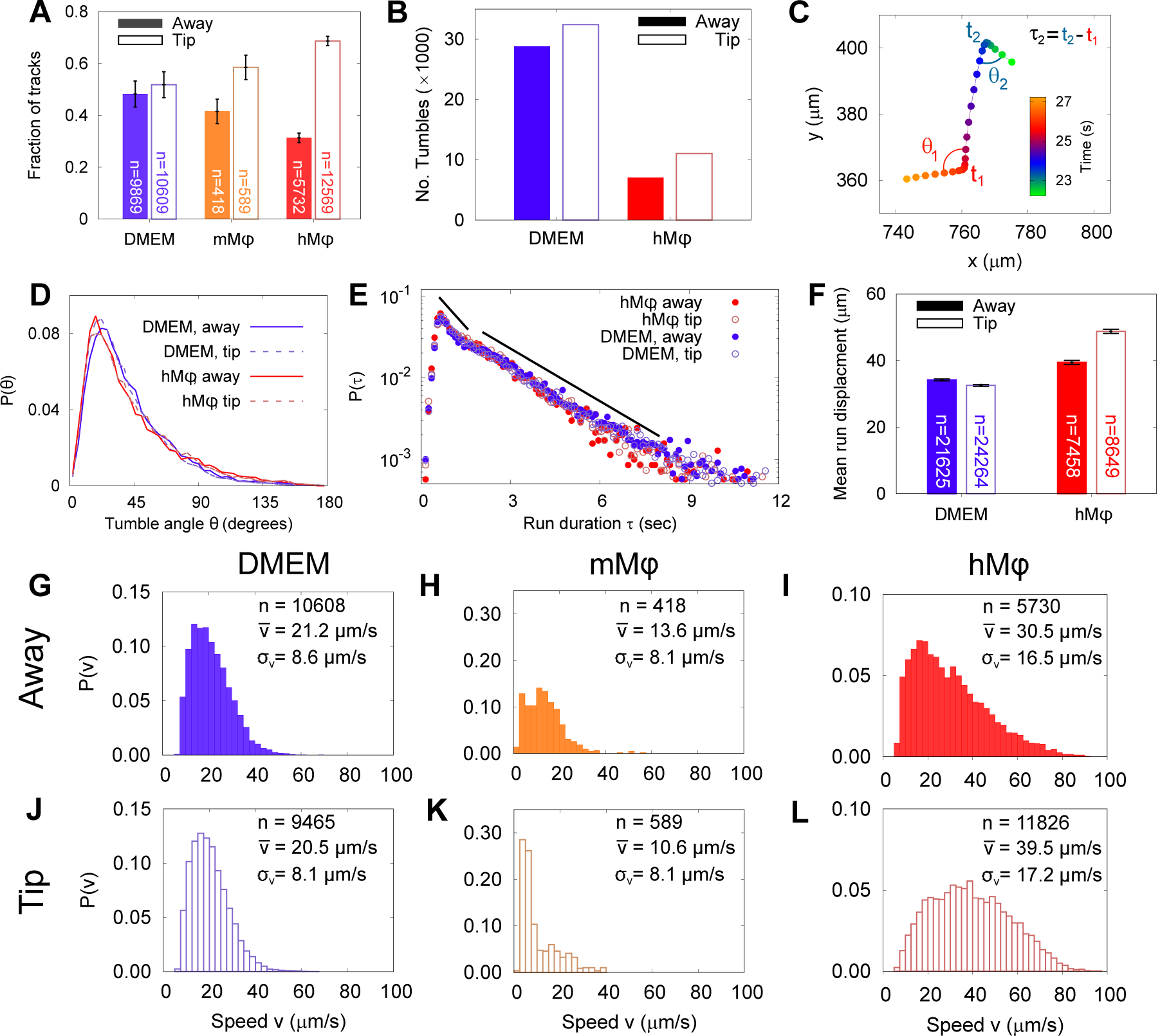
The run and tumble behaviour of META cells in the presence of an immunological stimulus. (A), Fractions of tracks from META cells promastigotes observed in a bulk suspension of cells distal (‘Away’, solid bars) and next to a loaded pipette tip immersed in the sample medium (‘Tip’, filled bars). The negative controls, in which tips were filled with DMEM agar, show negligible differences in the number of tracks proximal to the tip, versus the distal bulk. Pipette tips filled with either cultured J774.2 immortalised, mouse stomach-derived macrophages (mM*ϕ*) or fresh human blood-derived macrophages (hM*ϕ*) showed significant accumulations of cells around the pipette tips (error bars = SE). (B), The number of ‘tumble’ events for cells in control (DMEM) and test (hM*ϕ*) conditions. Exposure to hM*ϕ* reduces the number of tumble events considerably, consistent with a more persistent swimming direction. This effect impacts not only the cells that are in the immediate proximity of the human-derived macrophages (hM*ϕ*, Tip) but also those in the rest of the sample chamber. As the cells are free to explore the whole sample volume, any ‘activation’ of cells exposed to a macrophage stimulus may persist even when the cells themselves have left the immediate environment of the pipette tip. (C), x-y projection of a section of a META track, showing tumble times (*t*_1_, *t*_2_), reorientation angles (*θ*_1_, *θ*_2_) and intervening run duration (*τ*_2_). (D), The distribution of tumble angles remains unchanged in the presence of macrophage stimulus, suggesting an unbiased stochastic reorientation process. (E), Distribution of run durations. Contrary to what would be expected in the equivalent situation in chemotaxis in model bacterial systems, there is no increase in run duration in the presence of stimulus (hM*ϕ* ‘Tip’), compared to controls (hM*ϕ* ‘Away’, DMEM ‘Tip’, DMEM ‘Away’); the distribution of run durations is almost identical in all cases. (F), The mean displacement per run, showing an increase in the run displacement as a consequence of the faster run speed near the macrophage-loaded pipette tip. (error bars = SEM). (G-L), Normalised instantaneous cell speed distributions for cells distal versus proximal to stimulus. Control samples (DMEM; G,J) show minimal difference, but the presence of mouse- or human-derived macrophages causes a decrease or increase in swimming speed, respectively.

## Summary and Conclusions

*Leishmania* cells must survive changing environmental conditions throughout their lifecycle, and thus motile stages vary significantly in morphology and behavior. Pre-adaptation to the next host environment in parasitic protozoans has been well documented in trypanosomes [26, 24]. For *Leishmania*, this has been described in terms of changes in metabolism, gene expression [27], and the acquisition of a complement-resistant lipophosphoglycan coat [28, 29]. We find no bias in the chirality of swimming tracks in large populations. This leads us to conclude that helical swimming patterns resulting from broken structural symmetry are most likely introduced by small, random asymmetries in the cell body or position of the flagellum specific to individual cells. Our studies show that flagellar motility is a hitherto unrecognised pre-adaptation to infection of the human host. We show that run-and-tumble behaviour is a phenotypic marker of human-infective META stage parasites, and that this behaviour is suppressed in the presence of a chemotactic stimulus, causing cells to swim faster and straighter towards their target. We suggest that this provides an evolutionary advantage which promotes successful phagocytic uptake of non-replicative META cells, fundamental to human infection and parasite lifecycle progression. Through optimised flagellar motility and an as-yet uncharacterised sensing mechanism, *Leishmania* proactively drive their uptake by human immune cells.

### *Leishmania mexicana* cell culture

Promastigote parasites of *L.mexicana* (strain M379) were cultured in M199 and Grace’s media at 26^°^C as described previously [18]. All cells were maintained within low passage numbers (¡5) to ensure biological relevance to differentiation and infectivity was not compromised. PCF promastigotes were harvested mid-logarithmic phase at concentration 3–6×10^6^ cells/ml. This stock sample was diluted by a factor of 100 into fresh M199 medium to 5×10^4^ cells/ml for holographic tracking. To generate META promastigotes, PCF culture was passaged into Grace’s medium and cultured for seven days at 26^°^C [18, 30]. For META promastigote populations, cells were centrifuged in a 10% Ficoll gradient [31] to enrich for META cells as defined by morphology, molecular markers and macrophage infectivity. Purified META were transferred into fresh M199 medium and diluted to 5×10^4^ cells/ml for holographic tracking.

### Molecular validation of *L.mexicana* promastigote stages via qRTPCR

RNA was isolated from PCF and META populations concurrent to image capture, purified, reverse transcribed and analysed for *Histone H4* and *SHERP* mRNA levels relative to constitutive *nmt* levels as described previously [18].

### Macrophage cell culture

Immortalised mouse stomach macrophages (J774.2) and human blood-derived macrophages from resident hostages were isolated and cultured *in vitro* for 7 days in Dulbecco’s Modified Eagle Medium (DMEM) medium at 37^°^C prior to centrifugation. Approximately 1×10^4^ cells cells were resuspended in 10 *μ*L DMEM:1 % agar for each chemotactic assay sample. Macrophage viability was confirmed via alamar blue staining two hours after sample preparation.

### Sample chambers

To compare life cycle stages, glass sample chambers were constructed from slides and UV-curing glue, giving a final sample volume measuring approximately 20×15×1.2 mm^3^. These were loaded with *L.mexicana* in M199 and sealed with petroleum jelly before observation.

### Chemotaxis Assay

The stability of the chemotactic apparatus and resultant gradient was verified via a fluorescein-infused agar gradient establishment test (see Supp. Materials). The fluorescein-agar was then replaced in the 10 *μ*L pipette tip by different ‘Chemotactic bait’; including either DMEM:1 % agar alone, DMEM:1 % agar with immortalised mouse macrophages or with human blood-derived macrophages. Chemotaxis sample chambers were constructed as above with the addition of the 10 *μ*L pipette tip sealed into the sample chamber with petroleum jelly. The assembly was then placed on a temperature-controlled microscope stage at 34^°^C in a room set at 34^°^C to equilibrate for 30 minutes, allowing transient convection currents to dissipate and a chemical gradient to become established. The spatial profile of the putative chemical gradient was modelled using fluorescein in a separate experiment (see Supp. Inf.)

### Holographic video microscopy

The holographic microscopy setup was similar to that used previously [32, 33], with a few modifications. The samples were imaged on a Nikon Eclipse E600 upright microscope. The illumination source was a single-mode fibre-coupled laser diode with peak emission at 642 nm. The end of the fibre was mounted below the specimen stage using a custom adaptor and delivered a total of 15 mW of optical power to the sample. A Mikrotron MC-1362 monochrome camera was used to acquire videos of 3000 frames, at a frame rate of 50 Hz and with an exposure time of 100 *μ*s. A 10× magnification bright field lens with numerical aperture of 0.3 was used to acquire data at a video resolution of 1024×1024 pixels^2^, corresponding to a field of view measuring 1.44×1.44 mm^2^. The raw videos were saved as uncompressed, 8-bit AVI files.

### Holographic data reconstruction

From each individual video frame, we calculated a stack of 130 images, spaced at 10 *μ*m along the optical axis, sampling the optical field within a volume of 1.44×1.44×1 mm^3^. These sequences of images were calculated using the Rayleigh-Sommerfeld back-propagation scheme [34]. We localised the cells using a method based on the Gouy phase anomaly, as described in more detail elsewhere [21, 35]. This method segments features based on axial optical intensity gradients within a sample, allowing us to extract three-dimensional coordinates for individual cells in each frame. Lateral position uncertainties are approximately 0.5 *μ*m, while the axial performance is slightly worse at approximately 1.5 *μ*m. The latter is limited by the angular resolution of the microscope objective, and the scattering properties of the anisotropic cell bodies. A separate software routine was used to compare the putative cell coordinates extracted in each frame and associate those most likely to constitute cell tracks. The tracks were smoothed using piecewise cubic splines in order to remove noise in the cell coordinates and provide better estimates of cell velocity as described in previous work [33]. Examining the mean-squared displacement of the cells’ smoothed trajectories allowed us to discriminate between swimming and diffusing cells, and to discard the latter. The smoothing process also allowed for linear interpolation of missing data points, up to 5 points (equivalent to 0.1 second). Tracks with a duration shorter than three seconds were discarded.

### Frequency-domain analysis

To extract the flagellar beat and body roll frequencies, the smoothed trajectories were subtracted from the raw values, and the power spectrum of the residuals was calculated. The power spectra from all cells in the sample were summed to give the data in Fig. 1j. To illustrate the dominance of the two main frequency components, the track residuals were filtered in the frequency domain using a filter with two Gaussian pass bands with mean (standard deviation) frequencies of 1 Hz (1 Hz) and 20 Hz (2 Hz). The resulting signal contains the body roll and flagellar beat, removing extraneous noise, as shown in Fig. 1h.

### Track chirality analysis

The procyclic promastigote and metacyclic promastigote cell tracks were analysed to extract their chirality (handedness). This was done according to a scheme similar to that previously used for determining the chirality of flagellar waveforms [15]. We first constructed an array of displacement vectors **T**_*j*_ from a cell’s coordinates r_*j*_(*t*) by calculating **T**_*j*_ = **r**_*j*_ − **r**_*j*−1_, and obtained the local track handedness as the angle formed between vector **T**_*j*_ and the plane formed by the two previous segments (defined by **T**_*j*−2_ ^ **T**_*j*−2_), divided by the total contour length |**T**_*j*−2_| + |**T**_*j*−1_| + |T_*j*_|. We used this instead of (e.g.) the more conventional Frenet-Serret apparatus because the handedness (*H*) is mathematically well-behaved for straight trajectories.

### Run-tumble analysis

Tumbles were identified as large changes in swimming direction coupled with a drop in swimming speed. Quantitatively, we calculated

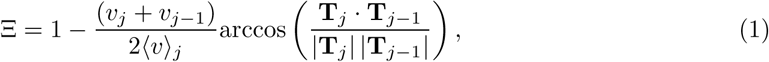

where we denote the local speed during segment **T**_*j*_ as *v*_*j*_ and the average speed of track *j* as ⟨ *v* ⟩ _*j*_. This quantity shows clear peaks when the cells change direction (‘tumbles’), as identified in Fig. 2d. Run duration *τ* was then defined as the time elapsed between two subsequent tumble events. The tumble events take a finite time, during which the cell is essentially stationary. We therefore define the tumble angle as the change in the cell’s swimming direction between two time points one second apart, with the tumble event halfway in between.

## Data availability

Data underlying the conclusions in this study are are available at the York Research Database (https://doi.org/10.15124/a8eabcd4-66c1-41a3-aa4b-423921b06568).

## Ethics Statement

Unrelated adults of both genders and diverse racial backgrounds local to Yorkshire, United Kingdom volunteered for tissue sampling for this research. After health screening by medical staff, informed consent was obtained from donors in compliance with ICH GCP, the UK Data Protection Act, and other regulatory requirements, as appropriate and under the University of York Department of Biology Ethics Committee (BEC)-approved study CL 201201 v2.3 to donate blood for a BEC-approved biomedical research under Project License PK 201702.

## Acknowledgements

This work was supported by Wellcome Trust grant nos. WT10472 and WT105502MA, EPSRC grant no. EP/N014731/1, and MRC grant no. MR/L00092X/1.

## Notes

### Competing Interest Statement

The authors have declared no competing interest.

